# Cross-transmission is not the source of new *Mycobacterium abscessus* infections in a multi-centre cohort of cystic fibrosis patients

**DOI:** 10.1101/582684

**Authors:** Ronan M. Doyle, Marc Rubio, Garth Dixon, John Hartley, Nigel Klein, Pere Coll, Kathryn A. Harris

## Abstract

**Background:** *Mycobacterium abscessus* is an extensively drug resistant pathogen that causes pulmonary disease particularly in cystic fibrosis (CF) patients. Identifying direct patient-to-patient transmission of *M. abscessus* is critically important in directing infection control policy for the management of risk in CF patients. A variety of clinical labs have used molecular epidemiology to investigate transmission. However there is still conflicting evidence as to how *M. abscessus* is acquired and whether cross-transmission occurs. Recently labs have applied whole-genome sequencing (WGS) to investigate this further and in this study we investigate whether WGS can reliably identify cross-transmission in *M. abscessus*.

**Methods:** We retrospectively sequenced the whole genomes of 145 *M. abscessus* isolates from 62 patients seen at four hospitals in two countries over 16 years.

**Results:** We have shown that a comparison of a fixed number of core single nucleotide variants (SNVs) alone cannot be used to infer cross-transmission in *M. abscessus* but does provide enough information to replace multiple existing molecular assays. We detected one episode of possible direct patient-to-patient transmission in a sibling pair. We found that patients acquired unique *M. abscessus* strains even after spending considerable time on the same wards with other *M. abscessus* positive patients.

**Conclusions:** This novel analysis has demonstrated that the majority of patients in this study have not acquired *M. abscessus* through direct patient-patient transmission or a common reservoir. Tracking transmission using WGS will only realise its full potential with proper environmental screening as well as patient sampling.

**Key points:**

- Whole genome sequencing should replace current molecular typing used routinely in clinical microbiology laboratories.
- Patient-to-patient spread of *M. abscessus* is not common.
- Environmental screening may provide a better understanding acquisition of *M. abscessus* infections.

## Background

*Mycobacterium abscessus* (recently renamed as *Mycobacteroides abscessus*) [1], is a group of three closely related subspecies *M. abscessus* subsp. *abscessus, M. abscessus* subsp. *massiliense* and *M. abscessus* subsp. *bolletii* [1,2]. These rapidly-growing, non-tuberculous mycobacteria cause chronic pulmonary disease, particularly in patients with cystic fibrosis (CF) and other chronic lung diseases. *M. abscessus* is an important pathogen that has emerged in the CF patient population that has been associated with poor clinical outcomes, especially following lung transplantation [3–5]. This is due, at least in part, to the extensive antibiotic resistance that makes infections with this organism difficult to treat [2,6]. CF patients infected with *M. abscessus* are frequently not listed for transplant, therefore the acquisition of this pathogen is considered to be a serious complication in this group.

The epidemiology of *M. abscessus* strains has been studied using Variable Nucleotide Tandem Repeats (VNTR) and Multi Locus Sequence Typing (MLST) [7]. The clustering of globally spread sequence types was confirmed with whole genome sequencing (WGS) and has provided greater resolution in how the various lineages are related as well as predicting possible transmission routes [8,9]. A dominant method of transmission of *M. abscessus* remains contested [10,11], with evidence for and against patient-to-patient transmission being the common route [8,12–14]. *M. abscessus* is ubiquitous in the environment with its niche hypothesised to be free-living amoeba [15,16], but due to the difficulties in isolating the organism, little has been done to track environment-to-patient acquisition. Confirmation of direct patient-to-patient transmission is important as it influences management of high-risk patients it could increase the effectiveness of infection control interventions by directing the use of limited resources.

In this retrospective study we assessed utility of using WGS to characterise subspecies, antimicrobial resistance (AMR) profiles and typing of *M. abscessus* isolates. We also wanted to utilise the data to investigate the scale of patient-to-patient transmission and whether identification of single nucleotide variants (SNVs) by WGS can confirm transmission. To do this we have sequenced the genomes 145 *M. abscessus* clinical isolates from a well characterised cohort of 62 patients from four hospitals in two countries over 16 years.

## Methods

### Patients and Samples collection

We collected 33 *M. abscessus* isolates from 30 patients at Hospital de la Santa Creu I Sant Pau (bcn_hsp), Hospital Clínic (bcn_hcl) and Hospital Vall d’Hebron (bcn_hvh), Barcelona, Spain and 112 isolates from 32 patients from Great Ormond Street Hospital (ldn_gos), London, UK (Supplementary table 1). Demographic and patient location data were obtained from the patient administration system and microbiological data from the laboratory information management system using SQL and Excel spreadsheets. Additional sources of information included CF and transplant databases. American Thoracic Society consensus guidelines were used to verify evidence of non-tubercuolous mycobacterial infection [17]. All investigations were performed in accordance with the Hospitals Research governance policies and procedures.

### DNA extraction and Whole-Genome Sequencing

One hundred and forty-five *M. abscessus* isolates from 62 patients were analysed using whole-genome sequencing. Briefly, DNA was extracted from all isolates as previously described [18] with some modifications: DNA was extracted from all isolates using Qiagen EZ1 Blood extraction kit with a previous step of bead-beating (Qiagen, Crawley, United Kindom). Then total DNA concentration was determined using a Qubit fluorometer (Thermofisher). Fifty nanograms of DNA was prepared using Nextera Library Preparation kit (Illumina) and post-PCR clean-up was carried out using Ampure XP beads (Beckman). Library size was validated using the Agilent 2200 TapeStation with Agilent D5000 ScreenTape System (Willoughby, Australia) and 150bp paired-end reads were sequenced on a NextSeq 550 system (Illumina). Raw sequencing reads have been deposited on ENA (study accession PRJEB31559).

### Multi Locus Sequence Typing (MLST) analysis

We used a custom bash script to extract the alleles of the multi-locus sequence typing (MLST) profile from the mapped reads to the reference genome. The MLST profile was obtained using the Institut Pasteur MLST database (http://bigsdb.pasteur.fr/mycoabscessus/mycoabscessus.html).

### Read mapping and variant calling

Sequenced reads for all samples were first mapped to *M. abscessus* subsp. *abscessus* ATCC 19977 using BBMap v37.90 (Joint Genome Institute). Single nucleotide variants (SNVs) were called against the reference genome using freebayes v1.2.0 [19] and variants were filtered to only include those at sites with a mapping quality >30, a base quality >30, at least five supporting reads, where the variant was present on at least two forward and reverse strand reads and present at the 5’ and 3’ end of at least two reads.

### Phylogenetic analysis

Potential regions of recombination were identified from the consensus genome sequences using Gubbins v2.3.1 [20]. Regions within the genome with low coverage (< 5x) were masked on a per sample basis and regions with low coverage across 75% of samples were masked across the entire dataset. A maximum likelihood tree was inferred from all samples using RAxML v8.2.4 [21] using a GTRCAT model with 99 bootstraps. Sub-species were identified for each sample based on their position upon this tree.

Separate sub-trees were also inferred for *M. abscessus* subsp. *massilense* sequences, as well as for *M. abscessus* subsp. *abscessus* ST-1 and ST-26 sequences. All samples in each sub-tree were mapped against a suitable reference. *M. abscessus* subsp. *massilense* str. GO 06 was used as the reference sequencing for study *massilense* sequences and the *de novo* assembly of the earliest ST-26 study sequence (ldn_gos_2_520) was used as a reference for other ST-26 samples. *M. abscessus* subsp. *abscessus* ATCC 19977 was again used as the reference for ST-1 sequences as it is the same sequence type. All sub-trees were generated using the same method outlined above, apart from ST-26 subtree, which did not use Gubbins but instead variants were filtered if 3 SNVs were found within a 100bp window.

### Sequence clusters

Sequence clusters to infer possible transmission were generated using three different methods on each subtree. First we used a SNV threshold that was based on the upper bounds of all within patient diversity applied to complete linkage hierarchical clustering based on pairwise SNV matrix. Secondly we assigned clusters using the R package rPinecone as it incorporates SNV thresholds and root-to-tip distances and so has been useful when applied to clonal populations [22]. Lastly we also used hierBAPS [23] to assign clusters, however due to the fact that all samples are included in the sequence clusters we found it was not appropriate for this study question. We made the assumption that any strains taken from different patients that were within sequence cluster constituted a possible transmission event.

### *De novo* assembly

All samples underwent *de novo* assembly of bacterial genomes using SPAdes and pilon wrapped in the Unicycler v0.4.4 package [24]. Assembled contigs were annotated using prokka v1.13 [25] and comparison of the accessory genome was generated using roary v3.12.0 [26]. To generate a list of genes that could be used to differentiate isolates we filtered the annotated genes to remove coding sequences (CDS) greater than 8000 bp and less than 250 bp, as well as those only present in a single sample and those present in every sample.

## Results

### *M. abscessus* population distribution

We obtained whole genome sequences for 145 *M. abscessus* isolates from 62 patients. Thirty-three *M. abscessus* from Barcelona subdivided into 24 *M. abscessus* subsp. *abscessus*, two *M. abscessus* subsp. *bolletii and* seven *M. abscessus* subsp. *massiliense*. A hundred and twelve *M. abscessus* from UK subdivided into 78 *M. abscessus* subsp. *abscessus,* one *M. abscessus* subsp. *bolletii and* 33 *M. abscessus* subsp. *massiliense*. Sample MLST definitions, VNTR and AMR associated mutations are shown in supplementary table 2.

### Possible transmission within *M. abscessus* clusters

To confirm possible transmission between patients we required their isolate genomes to be clustered together by two independent methods and epidemiological evidence that both patients were at the same hospital during the same time period. Using WGS data we inferred a phylogenetic tree from reference genome SNV matrix for all patients (Figure 1). We observed two low variant clusters of isolates that corresponded to ST-1 and ST-26 Pasteur MLST profiles (VNTR II and I respectively), as well as other closely related *M. abscessus susp. massilense* isolates between patients. We used a SNV matrix from mapping against a reference (*M. abscessus subsp. abscessus* ATCC19977), as well as hierBAPS and rPinecone to predict sequence clusters. The sequence clusters generated from the single reference SNV matrix provided no further information than the MLST profiles, and in many cases provided spurious findings with large groups of isolates clustered with no epidemiological link (Supplementary Figure 1). This included large sequence clusters relating to a single MLST type which included isolates from different hospitals and countries.

**Figure 1.**
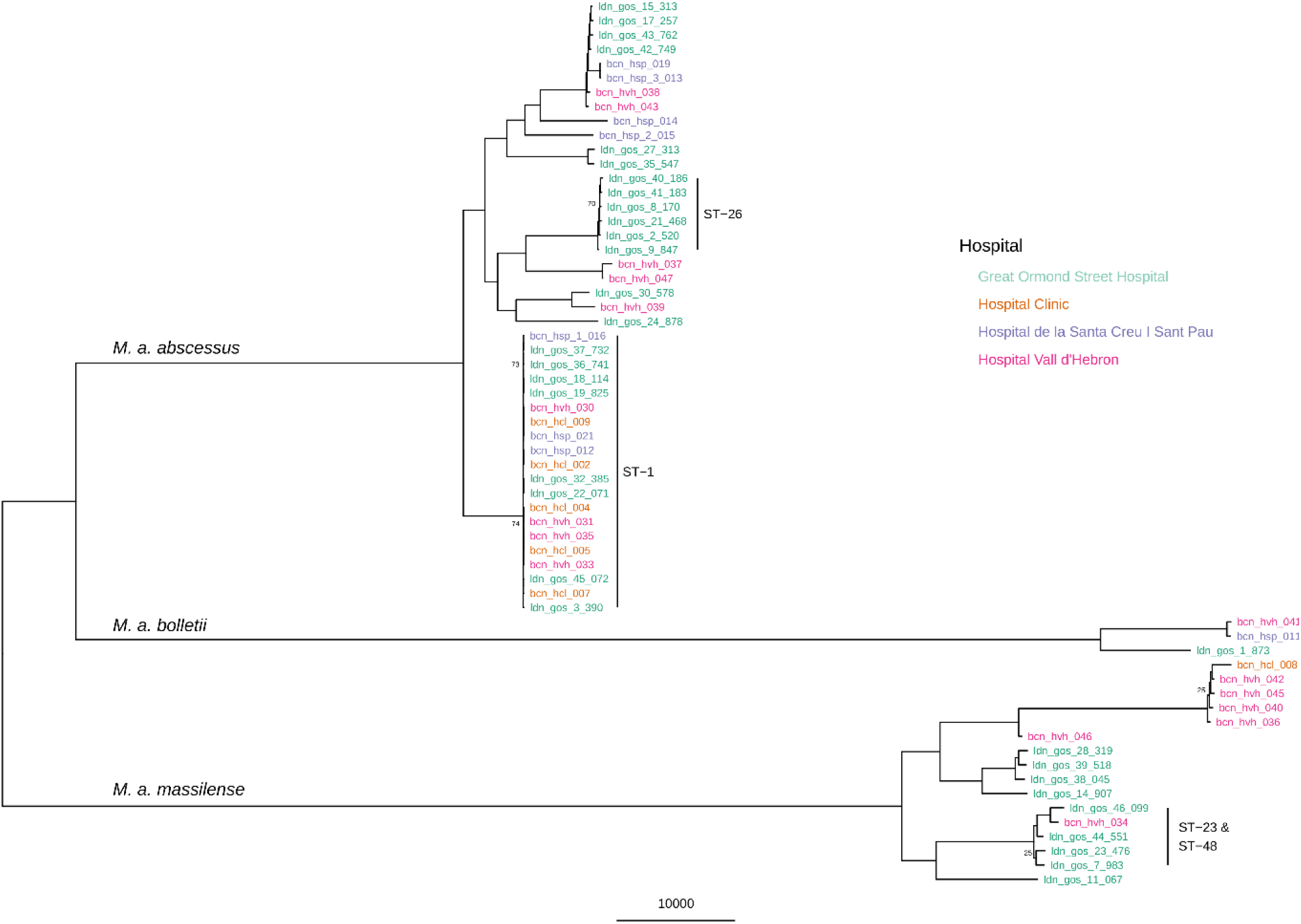
Maximum likelihood single nucleotide variant (SNV) tree using only the earliest isolated sample from all 62 patients. SNVs were identified from mapping reads to ATCC19977 *M. abscessus subsp. abscessus* reference genome. Sample names are highlighted in colour based on what hospital they were isolated from: Great Ormond Street Hospital, London, UK, Hospital Clínic, Barcelona, Spain, Hospital de la Santa Creu i Sant Pau, Barcelona, Spain, and Hospital Vall d’Hebron, Barcelona, Spain. The scale bar represents the number of single nucleotide variants and node bootstrap scores below are shown if below 75.

Mapping to a single reference genome led to the inability of a single SNV cut-off, or model, to exclude unrelated isolates from sequence clusters because the number of pairwise SNV distances varied greatly between both subspecies and specific lineages which (Figure 2). For example, the pairwise median (interquartile range) SNV distance between just ST-1 isolates was 73 (62 – 81) compared to 29589 (27701 – 63703) for all *M. abscessus subsp. abscessus* isolates. The same differences were seen in *M. abscessus subsp. massilense* as well with a pairwise median (IQR) SNV distance between ST-23 and ST-48 isolates of 2084 (960 – 7274) compared to 70545 (59947 – 71891) across all isolates from the subspecies.

**Figure 2.**
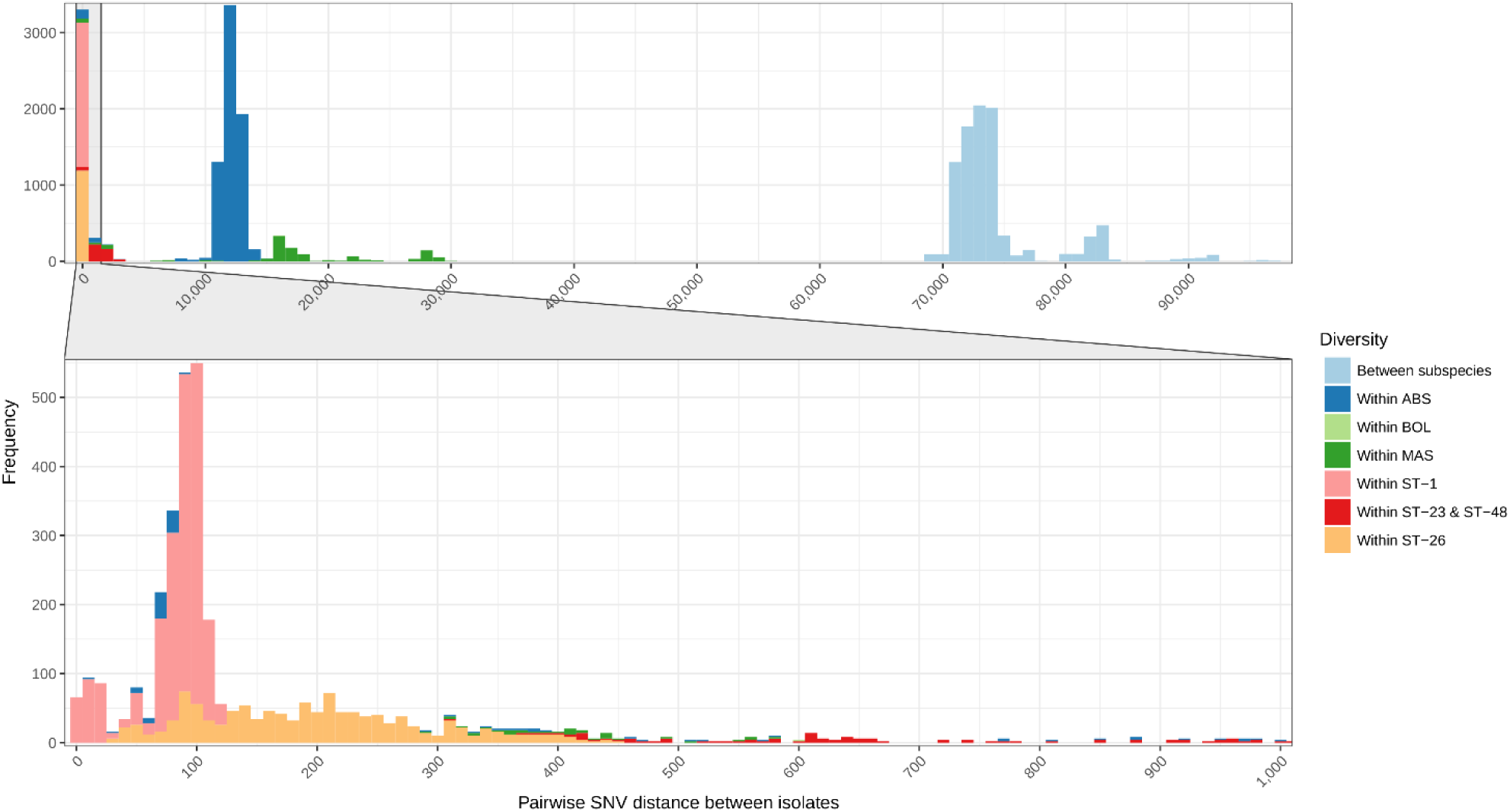
Frequency of pairwise single nucleotide variant (SNV) distances between all isolates. SNVs were identified from mapping sequence reads to *M. abscessus* subsp. *abscessus* ATCC19977. The full plot includes all samples while the bottom subsidiary plot only includes isolates that have a pairwise difference between zero and 1000 SNVs.

### Sub-tree sequence clusters

The variation in the scale of diversity within subspecies and sequence type hampered efforts to capture possible transmission events. In order to improve accuracy of sequence clustering, multiple sub-trees were made for closely related isolates using a more suitable reference sequence. We separated *M. abscessus subsp. abscessus* and *M. abscessus subsp. massilense* isolates, as well as further sub-trees for ST-1 (VNTR II), ST-26 (VNTR I) and ST-23/ST-48 (VNTR III) isolates. We also integrated the presence of accessory genes when interrogating possible sequence clusters for transmission (Figures 3, 4 & 5). Sequence clusters were assigned for each sub-tree using both a single SNV threshold (Supplementary Figure 2) and rPinecone. Overall we found that predicting transmission from the sub-trees reduced the number of different patients clustered together from 46 to 19 and the number of possible sequence clusters suggesting patient-to-patient transmission from 11 to seven.

**Figure 3.**
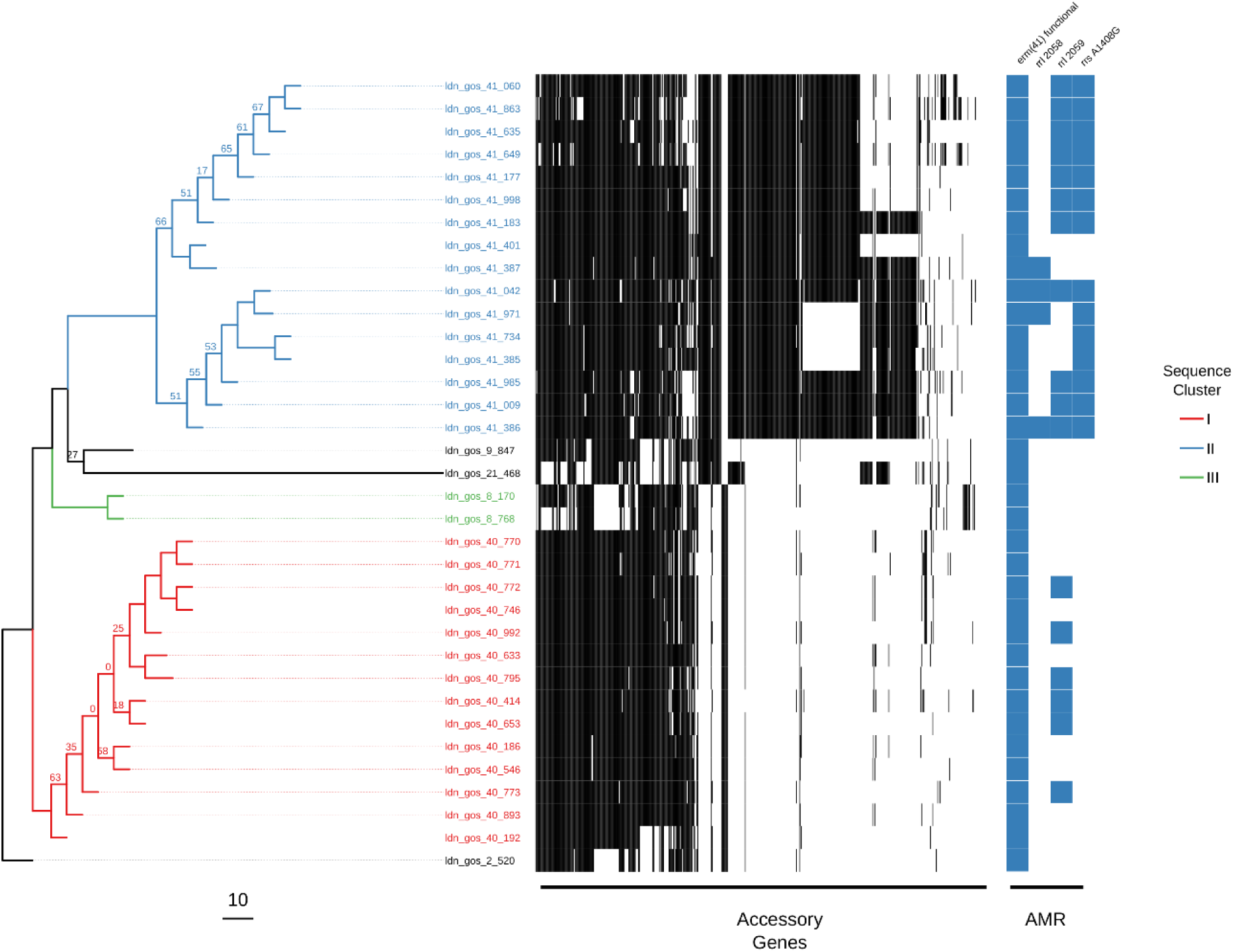
Maximum likelihood single nucleotide variant (SNV) tree for all ST-26 isolates. SNVs were identified from mapping reads to a de-novo assembled study isolate genome (ldn_gos_2_520). Samples are highlighted based on inclusion in sequence clusters. The tree is annotated with the presence (black) and absence (white) of accessory genes as well as the presence of AMR associated genes and mutations. This included presence of a functional *erm(41)* gene conferring inducible resistance to macrolides, presence of two *rrl* mutations conferring high level macrolide resistance and the presence of mutation in *rrs* conferring high level amikacin resistance. The scale bar represents the number of single nucleotide variants and node bootstrap scores below are shown if below 75.

**Figure 4.**
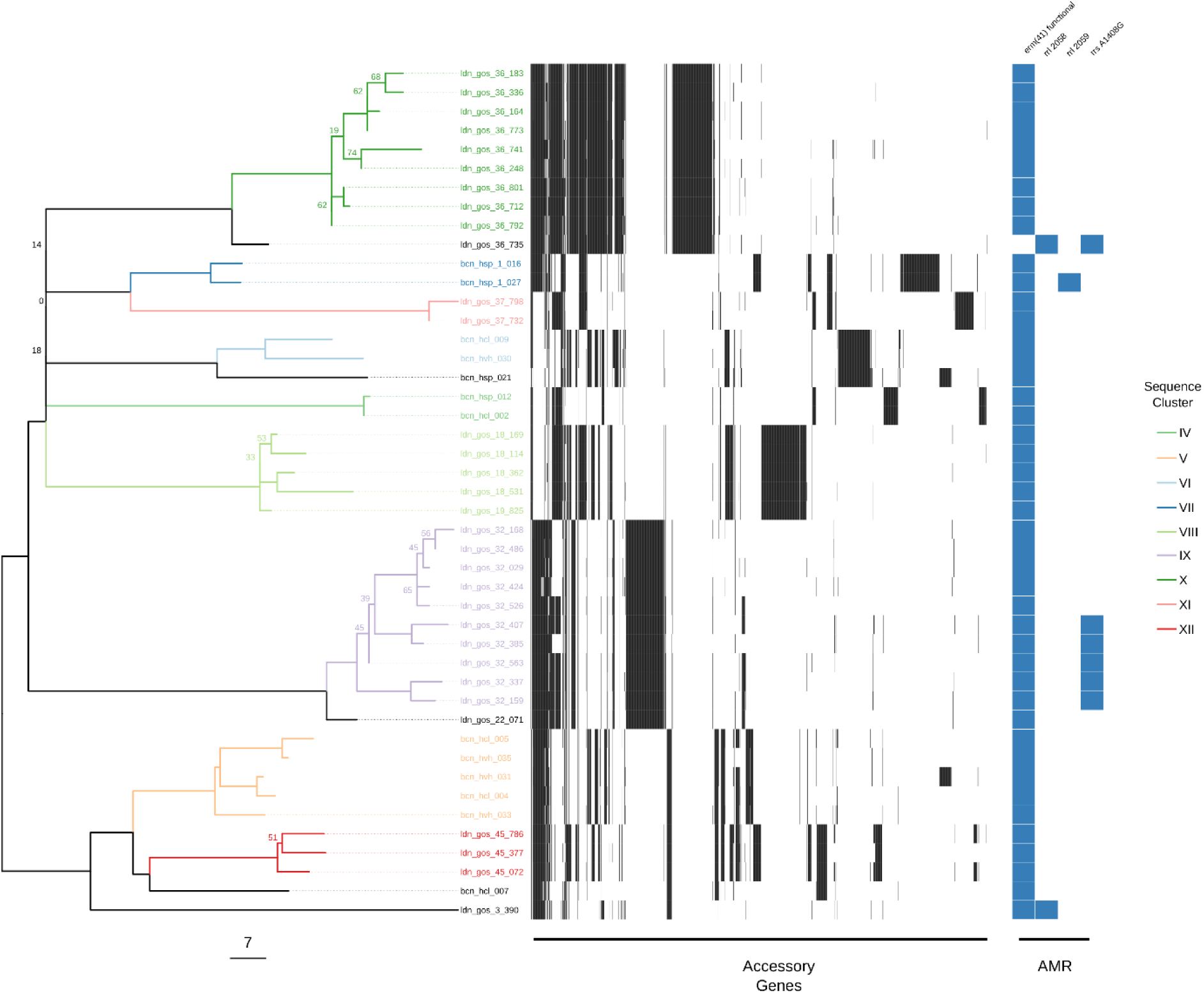
Maximum likelihood single nucleotide variant (SNV) tree for all ST-1 isolates. SNVs were identified from mapping reads to *M. abscessus* subsp. *abscessus* ATCC19977. Samples are highlighted based on inclusion in sequence clusters. The tree is annotated with the presence (black) and absence (white) of accessory genes as well as the presence of AMR associated genes and mutations. This included presence of a functional *erm(41)* gene conferring inducible resistance to macrolides, presence of two *rrl* mutations conferring high level macrolide resistance and the presence of mutation in *rrs* conferring high level amikacin resistance. The scale bar represents the number of single nucleotide variants and node bootstrap scores below are shown if below 75.

A total of 18 sequence clusters (I – XVIII) were identified (listed in supplementary table 2), 15 of these were within the sub-trees (I – XV), and seven clusters contained samples from more than one patient (IV, V, VI, VIII, XIV, XVI & XVII). All sequence clusters contained isolates from a single country with no evidence of international transmission. We found no evidence of transmission between patients within ST-26. (Figure 3). Within ST-1, four clusters (IV, V, VI and VIII) containing samples from more than one patient were found. Three of these clusters (IV, V and VI) contained isolates from nine patients from multiple hospitals within Barcelona. Only two of these patients were in hospital during the same time period (cluster VI: bcn_hcl_009 and bcn_hvh_30), but both were treated in different hospitals. Cluster VIII suggested transmission between two patients (ldn_gos_18 and ldn_gos_19) who were siblings with previously assumed either direct transmission or common reservoir [13] (Figure 4). A single cluster (XIV) containing samples from two patients (ldn_gos_46 and ldn_gos_7) was found among ST-23 isolates. However the two strains were isolated from samples taken nine years apart (Figure 5). Patient ldn_gos_7 was already positive for *M. abscessus* on first admission to GOSH, and the two patients were present at the lung function lab within a month of each other on two occasions, but never in the same location at the same day, and never admitted to the same ward.

**Figure 5.**
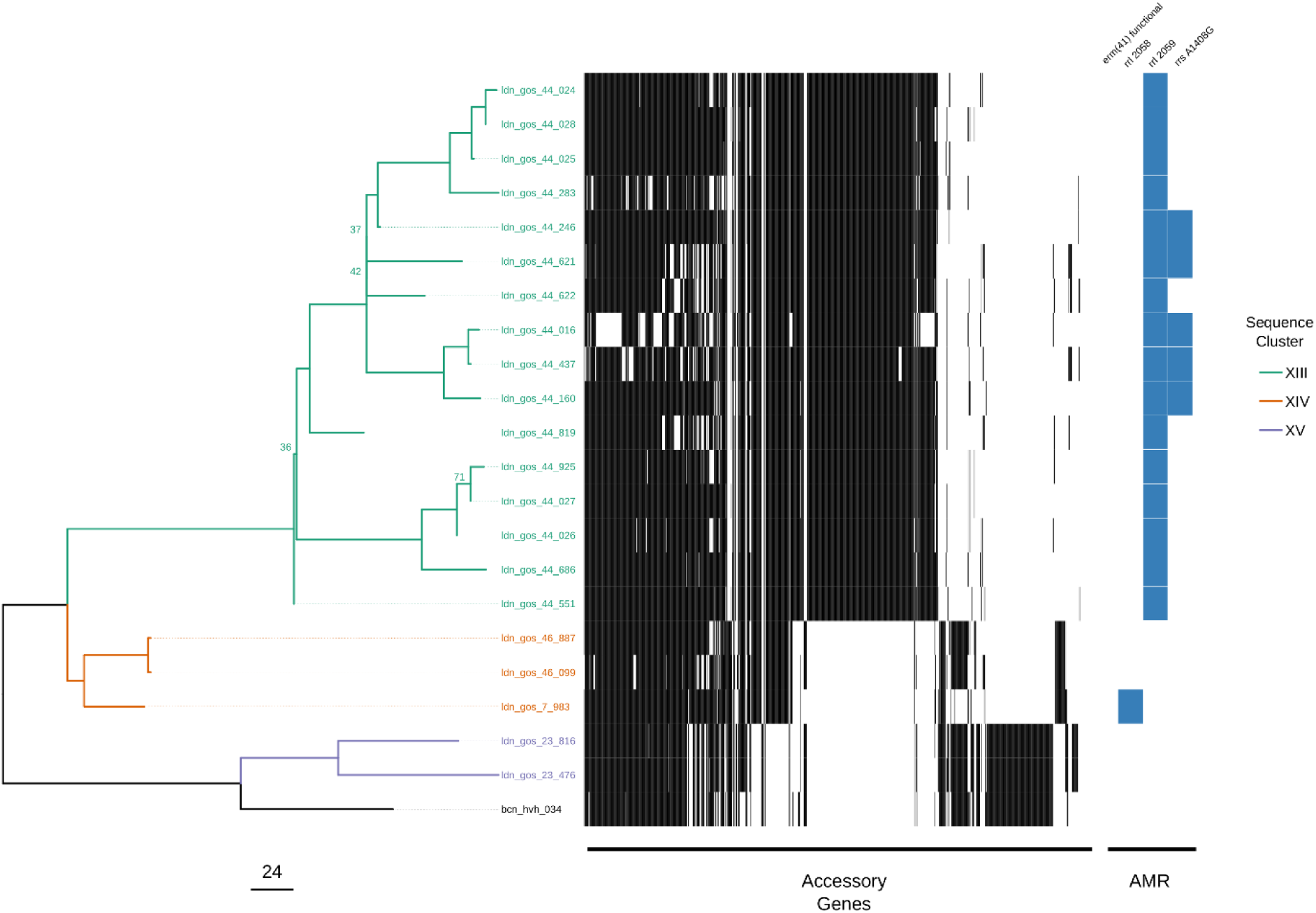
Maximum likelihood single nucleotide variant (SNV) tree for all ST-23 and ST-48 isolates. SNVs were identified from mapping reads to *M. abscessus* subsp. *massilense* GO 06. Samples are highlighted based on inclusion in sequence clusters. The tree is annotated with the presence (black) and absence (white) of accessory genes as well as the presence of AMR associated genes and mutations. This included presence of a functional *erm(41)* gene conferring inducible resistance to macrolides, presence of two *rrl* mutations conferring high level macrolide resistance and the presence of mutation in *rrs* conferring high level amikacin resistance. The scale bar represents the number of single nucleotide variants and node bootstrap scores below are shown if below 75.

All samples found within their respective clusters also contained similar accessory gene profiles with the median (IQR) shared percentage of accessory genes within a sequence cluster being 89% (79% – 94%) compared to 18% (12% – 37%) for isolates not in the same sequence cluster.

For the 32 GOSH CF patients included in the study, 16 became infected with *M. abscessus* after their first visit to clinic (Supplementary Table 1), however transmission confirmed by both WGS and epidemiological data could only be identified in one case (gos_19) thus suggesting a different route of acquisition for the rest of these patients.

## Discussion

This study has shown that whole genome sequencing of *M. abscessus* isolates can determine sub-species, identify previously reported AMR associated mutations and provide common typing definitions in a single workflow. This single method can replace the multiple existing molecular assays used in clinical microbiology laboratories to provide the same information and could be used to predict novel resistance variants [27]. We used the WGS data to investigate the likelihood of cross-transmission and found 43 (69%) patients had unique isolates that did not cluster with other patients. We identified seven sequence clusters from the remaining 19 patients but only one pair of patients (ldn_gos_18 and ldn_gos_19) had a plausible epidemiological link to support possible patient-to-patient transmission, as they were siblings. All other patients with genetically similar strains were either isolated in different countries, different hospitals or isolated from samples that were taken years apart, making direct transmission of these strains extremely unlikely.

Every *M. abscessus* isolated from a GOSH patient was sequenced and so the dataset generated represents a complete picture of *M. abscessus* infection in this hospital, which is vital for inferring transmission. Most of these patients were only attending clinics at GOSH, therefore this study has captured all of their *M. abscessus* isolates and they are unlikely to have been in contact with *M. abscessus* positive patients at other hospitals (Supplementary table 1). Therefore, if direct patient-patient transmission was occurring frequently we would expect to see evidence of it here. In contrast to this we found that the majority of patients in this study had unique strains and the majority of sequence clusters were multiple isolates from the same patients. This study confirms previous findings that despite many *M. abscessus* negative patients spending considerable time on the same wards as patients with ongoing *M. abscessus* infections they did not subsequently acquire genetically similar isolates.

We have therefore found that a fixed number of SNVs cannot be reliably used to infer cross-transmission across all *M. abscessus* isolates as there seems to be irreconcilable differences in the substitution rate between both sub-species and dominant clones. These difficulties are similar to those seen in *Legionella pneumophila* outbreaks where the majority of cases can belong to only a few sequence types [26]. *L. pneumophila* can also display different scales of genetic diversity within different sequence or genotypes and so it is also recognised that a single SNV threshold cut-off will not provide sufficient discriminatory power [27]. When using WGS to infer relatedness in *M. abscessus* there has previously been an attempt to find an absolute threshold which can rule in or rule out strains into a transmission event. This has previously been placed as below 25-30 SNVs [8,14,28,29]. From our findings we would advocate using a suitable genetically similar reference sequence when carrying out core genome SNV calling, especially for the dominant clones such as ST-1 and ST-26. There is a large amount of variation within the genomes of *M. abscessus* [30] and so the use of a single reference such as *M. abscessus subsp. abscessus* ATCC 19977 will mask many differences between strains and generate spurious clusters of genetically similar sequences. Where a suitable reference is not available we recommend using a high quality draft de-novo assembly of the first isolated sample to compare other isolates against as in the example of the ST-26 samples in this study (Figure 3).

In addition to core genome SNV analysis we have also found the integration of accessory genome information is a useful indicator of relatedness within *M. abscessus* isolates that can be used to further interrogate assigned sequence clusters. Generally there was good concordance between the proportion of putative genes shared and the SNV distance between two samples. This is helped by using a closely related reference sequences to map sequence reads against. We have seen in this study, and previously [31], diversity in the accessory genome profiles as well as in the number of SNPs and AMR associated mutations taken from multiple samples from the same patient on the same day. However we have always found inter-patient diversity to be greater than that seen within the same patient. This would suggest that any direct transmission between patients of even minority populations would still be identified by WGS and, taken together, the data suggests that person-to-person transmission of *M. abscessus* in paediatric patients in our institution is very uncommon. In this study we have an example of two patients with transmission predicted by genomic epidemiology (ldn_gos_7 and ldn_gos_46) that had attended a lung function laboratory on three occasions within a month of each other. In this case, the only way transmission could have occurred is if ldn_gos_7 who was already infected contaminated the environment and this then transmitted to ldn_gos_46. The predominant view [8] that human-to-human transmission occurs via contamination of fomites by respiratory secretions could explain this, although no other instances of this appeared to have occurred, despite numerous other CF patients attending the unit over many years. What is harder to explain is that for this to be the case, the interval between exposure and culture positivity was nine years. It could be that *M. abscessus* remains present but undetectable by conventional methods for this time period, or intriguingly could cause latent infection, like what occurs with *Mycobacterium tuberculosis*. To the best of our knowledge, this has never been a demonstrated part of the pathogenesis of *M. abscessus* infection, and maybe worthy of further investigation.

In agreement with previous studies we have found an international distribution of *M. abscessus* dominant clones [8]. We have found WGS to be useful to confirm whether different patient’s strains are unrelated, even within the dominant clones, but it has been far more difficult to reach definite conclusions about cross-transmission. Without environmental samples we cannot rule out the possibility of intermediate sources of infection and so WGS as a tool for tracking cross-transmission in *M. abscessus* will only realise its full potential with proper screening of environmental sources alongside longitudinal patient sampling.

## Funding

This work was supported by the National Institute for Health Research; EMBO Short-Term Fellowship [7307 to M.R.] and the European Association of National Metrology Institutes [15HLT07 to R.D.]

## Acknowledgements

We thank the Biomedical Scientist team for sample collection at Great Ormond Street Hospital as well as Dr Julià Gonzalez and Dr Teresa Tórtola for sample collection at Hospital Clinic and Hospital de la Vall d’Hebron, respectively.

## Supplementary material

**Supplementary table 1. Study patient information.**

**Supplementary table 2. Information on all individual *M. abscessus* isolates included in this study.**

**Supplementary Figure 1. Maximum likelihood single nucleotide variant (SNV) tree for all isolates in this study.** The tree is annotated with sequence clusters that are defined either by (from left-to-right) MLST, SNV threshold, hierBAPS and rPinecone as well as the presence of AMR associated gene and mutations. This included presence of a functional *erm(41)* gene conferring inducible resistance to macrolides, presence of two *rrl* mutations conferring high level macrolide resistance and the presence of mutation in *rrs* conferring high level amikacin resistance. The scale bar represents the number of single nucleotide variants and node bootstrap scores below are shown if below 75.

**Supplementary Figure 2. Frequency of pairwise single nucleotide variant (SNV) distances between samples after sub-tree analysis**. Figure 3A shows pairwise differences from the ST-1 subtree. Figure 3B shows pairwise differences from the ST-26 subtree. Figure 3C shows pairwise differences from the ST-23 and ST-48 subtree.

