## Supplementary Material for "Cross-transmission is not the source of new *Mycobacterium abscessus* infections in a multi-centre cohort of cystic fibrosis patients"

**Supplementary table 1. Study patient information.**

**Supplementary table 2. Information on all individual *M. abscessus* isolates included in this study.**

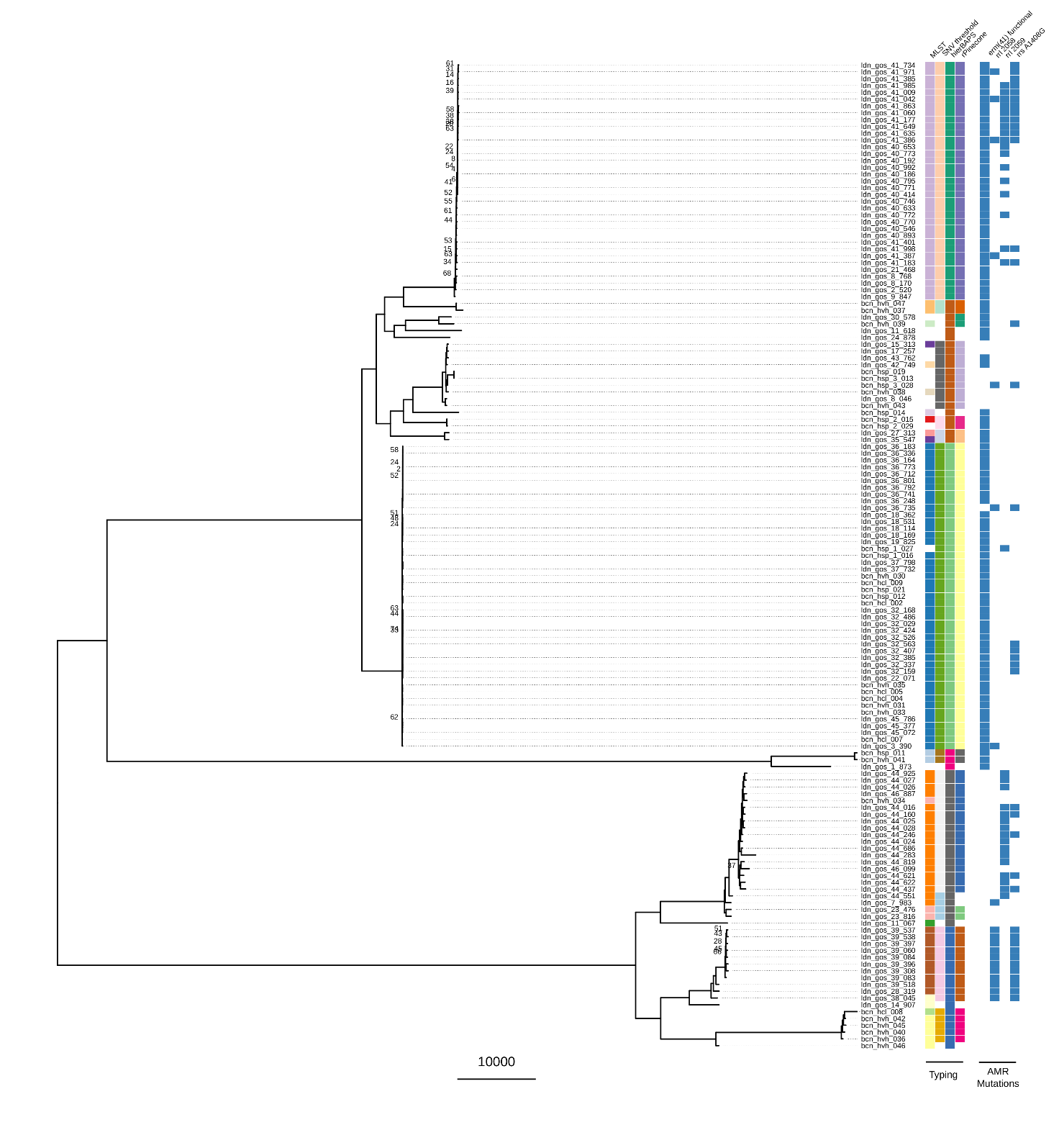

**Supplementary Figure 1.** **Maximum likelihood single nucleotide variant (SNV) tree for all isolates in this study.** The tree is annotated with sequence clusters that are defined either by (from left-to-right) MLST, SNV threshold, hierBAPS and rPinecone as well as the presence of AMR associated gene and mutations. This included presence of a functional *erm(41)* gene conferring inducible resistance to macrolides, presence of two *rrl* mutations conferring high level macrolide resistance and the presence of mutation in *rrs* conferring high level amikacin resistance. The scale bar represents the number of single nucleotide variants and node bootstrap scores below are shown if below 75.

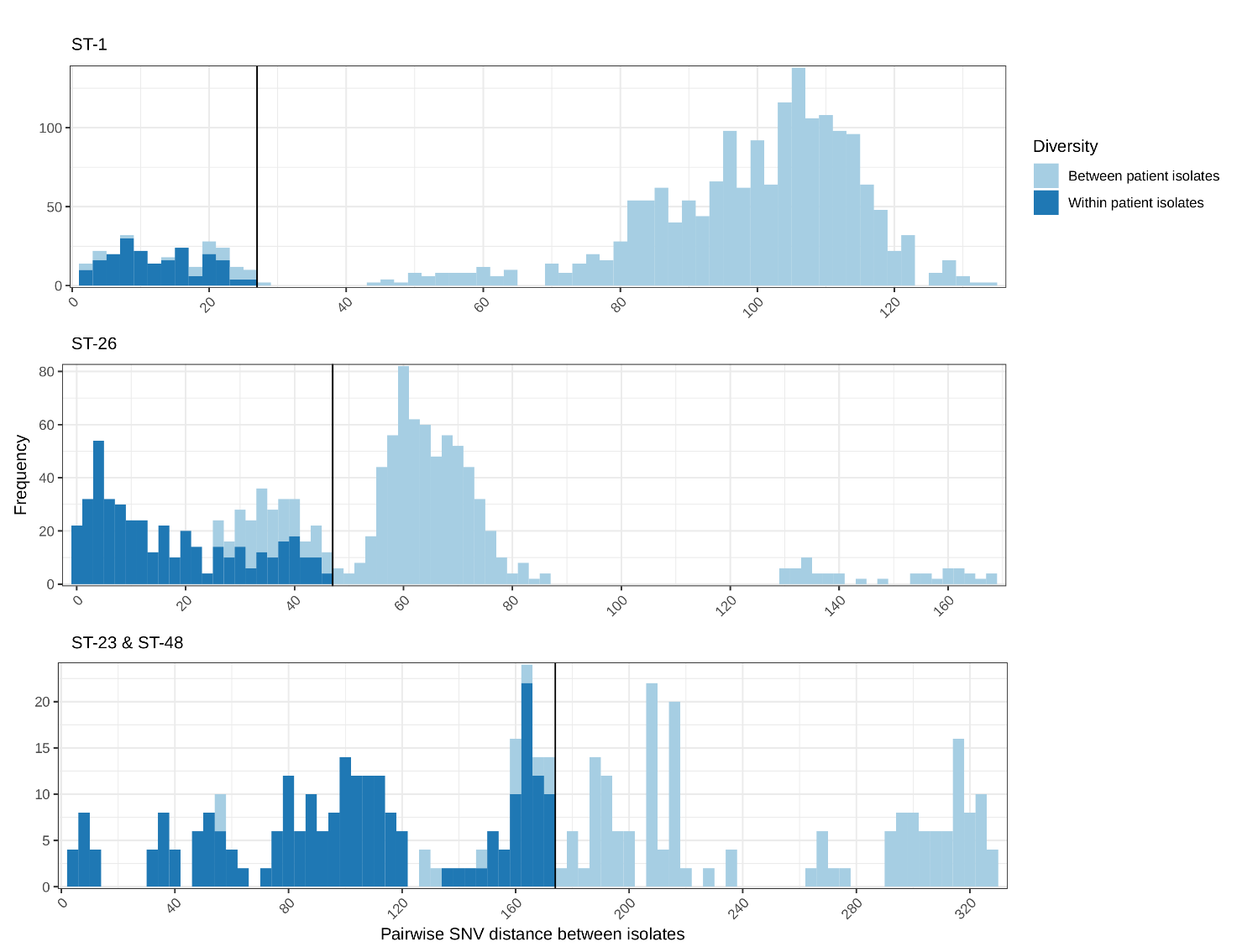
**Supplementary Figure 2. Frequency of pairwise single nucleotide variant (SNV) distances between samples after sub-tree analysis**. Figure 3A shows pairwise differences from the ST-1 subtree. Figure 3B shows pairwise differences from the ST-26 subtree. Figure 3C shows pairwise differences from the ST-23 and ST-48 subtree.
